# Detection of EGFR deletion using unique RepSeq technology

**DOI:** 10.1101/584045

**Authors:** Thuraiayah Vinayagamoorthy, Dahui Qin, Fei Ye, Minghao Zhong

## Abstract

We are reporting a novel sequencing technology, RepSeq (Repetitive Sequence), that has high sensitivity, specificity and quick turn-around time. This new sequencing technology is developed by modifying traditional Sanger sequencing technology in several aspects. The first, a homopolymer tail is added to the PCR primer(s), which makes interpreting electropherograms a lot easier than that in traditional Sanger sequencing. The second, an indicator nucleotide is added at the 5’end of the homopolymer tail. In the presence of a deletion, the position of the indicator nucleotide in relation to the wild type confirms the deletion. At the same time, the indicator of the wild type serves as the internal control. Furthermore, the specific design of the PCR and/or sequencing primers will specifically enrich/select mutant alleles, which increases sensitivity and specificity significantly. Based on serial dilution studies, the analytical lower limit of detection was 1.47 copies. A total of 89 samples were tested for EGFR exon 19 deletion, of which 21 were normal blood samples and 68 were samples previously tested by either pyrosequencing or TruSeq Next Generation Sequencing Cancer Panel. There was 100 % concordance among all the samples tested. RepSeq technology has overcome the shortcomings of Sanger sequencing and offers an easy-to-use novel sequencing method for personalized precision medicine.

## Introduction

The detection of somatic mutations in the epidermal growth factor receptor (EGFR) is the key for choosing first line targeted therapies for treating patients with late stage non-small cell lung cancer (NSCLC) (1,2,3). EGFR deletion mutations constitute a key component for the first line of targeted chemotherapy. These deletions are mostly confined to exon 19, from 2230 nt to 2260 nt of the reading frame that corresponds to amino acid changes in the cytoplasmic domain of the EGFR protein. Based on sample matrix, detection of EGFR deletions imposes five challenges; being a hyper variable region, the deletion could be anywhere in the above-mentioned region, there could be multiple deletions of varying number of nucleotides on the same allele, deletion could be homozygous or heterozygous, the copies of the deletion alleles could be low, and finally, the mutant allele is usually mixed wild type allele at different ratios. RepSeq technology has been developed to address most of these challenges. We have selected the most common exon 19 deletion, EGFR L747-A750, that offers the choice of treatment with Afatinib, Gefitinib, or Erlotinib (4) as an example to illustrate RepSeq technology. There are several FDA approved tests and a variety of laboratory developed tests (LDTs) to detect EGFR exon 19 deletion from clinical samples (5,6). All these tests are based on one of three major platforms; endpoint PCR, real-time PCR, and sequencing. The sequencing platforms generate a nucleotide sequence, and are hence considered an accurate confirmation of mutation detection. There are three commonly used sequencing platforms: Sanger sequencing, pyrosequencing and next generation sequencing (NGS). Sanger sequencing is the reference method for detecting EGFR L747_A750 deletion, with a deletion being determined by an overlap of the electropherogram sequences from the deletion and the wild type (7). Such an overlap generates scrambled nucleotide sequences that can be difficult to decipher to make the call on the nucleotide sequence. As an alternative, pyrosequencing is used routinely to detect EGFR L747_A750 deletion from FFPE samples (8). The nucleotide read outs from pyrosequencing require experience to call the results with confidence. Unlike germline mutations, where the copy number is high, EGFR L747_A750 is usually a somatic mutation, and different samples have different mutant to wild type ratios, adding another level of complexity to detect mutants in the presence of a n abundance o f wild type EGFR. To overcome the challenges with Sanger sequencing in determining a true deletion, and to increase the sensitivity to detect the EGFR L747_A750 somatic mutation, a new platform technology, Repseq, was developed. This manuscript presents the evaluation of the EGFR L747_A750 detection from FFPE samples and provides a comparison to pyrosequencing and TruSeq cancer panel (Illumina, USA).

## MATERIALS AND METHODS

### RepSeq platform technology

The RepSeq process includes extraction of total DNA from FFPE samples, followed by amplicon sequencing and analysis by capillary electrophoresis. If the sample carries a somatic deletion, there will be two amplicons generated, one carrying the mutant with deletion region, and the other carrying the wild type. Purified PCR products will be simultaneously sequenced using both wild type and mutant (deletion)-specific sequencing primers. Positioning of primers for amplification and sequencing is shown (Figure 1A & 1B). In order to distinguish the deletion from that of the wild type, RepSeq carries two modifications to Sanger sequencing: (a) the lower PCR primer carries a three nucleotide (Adenosine-Thymidine-Thymidine) repetitive sequence with a guanidine nucleotide at its 5’ end as an indicator of one end of the PCR product. (b) Two types of sequencing primers are used, the wild type sequencing primers and mutant primer. The wild type sequencing primers consist of two designs: selective and non-selective. The non-selective wild type sequencing primer anneals to the nucleotide sequence upstream of the deletions (Figure 1A). The selective wild type sequencing primer, however, not only anneals to the nucleotide sequence upstream of a deletion, but also anneals to the nucleotide sequence with in the deletion region (Figure 1B). The mutant (deletion) sequencing primer has only one design, which primes across the deletion region (Figure 1B). In the absence of a deletion, the sequencing result will show one wild type nucleotide sequence ending with a cysteine. If there is a deletion, the sequencing result will show two nucleotide sequences, one generated by the wild type sequencing primer ending with a cysteine, and a shorter nucleotide sequence generated by mutant sequencing primer also ending with a cysteine. There are a number of different deletions in EGFR. If the copies of the deletion alleles in the sample is more than 50%, a single sequencing primer (Non-selective) that anneals upstream of the deletion could be used to detect both the deletions and the wildtype. Depending on the number of deletions in the sample, a proportional number of deletion indicator signals will appear in the detection region. However, if specific deletions have low copies in the samples, one could use deletion specific sequencing primers that span across the specific deletion regions to increase the test sensitivity. These sequencing primers have different priming sites. Although at the proximal end of the sequences will not be distinguishable, at the distal end, the ‘C’ signal from the mutant will be among the TAA repeats, and hence could be detected.

**Figure 1.**
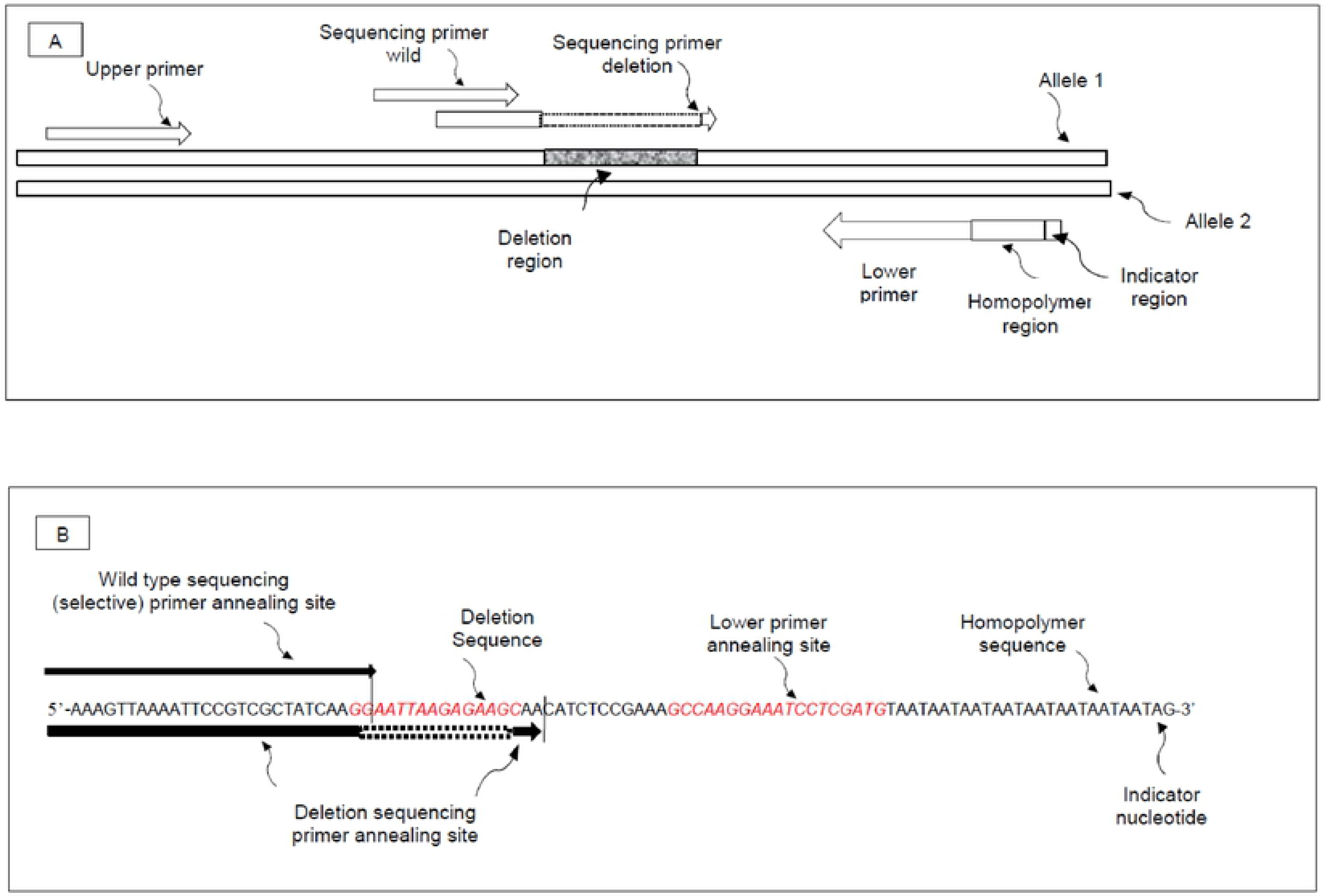
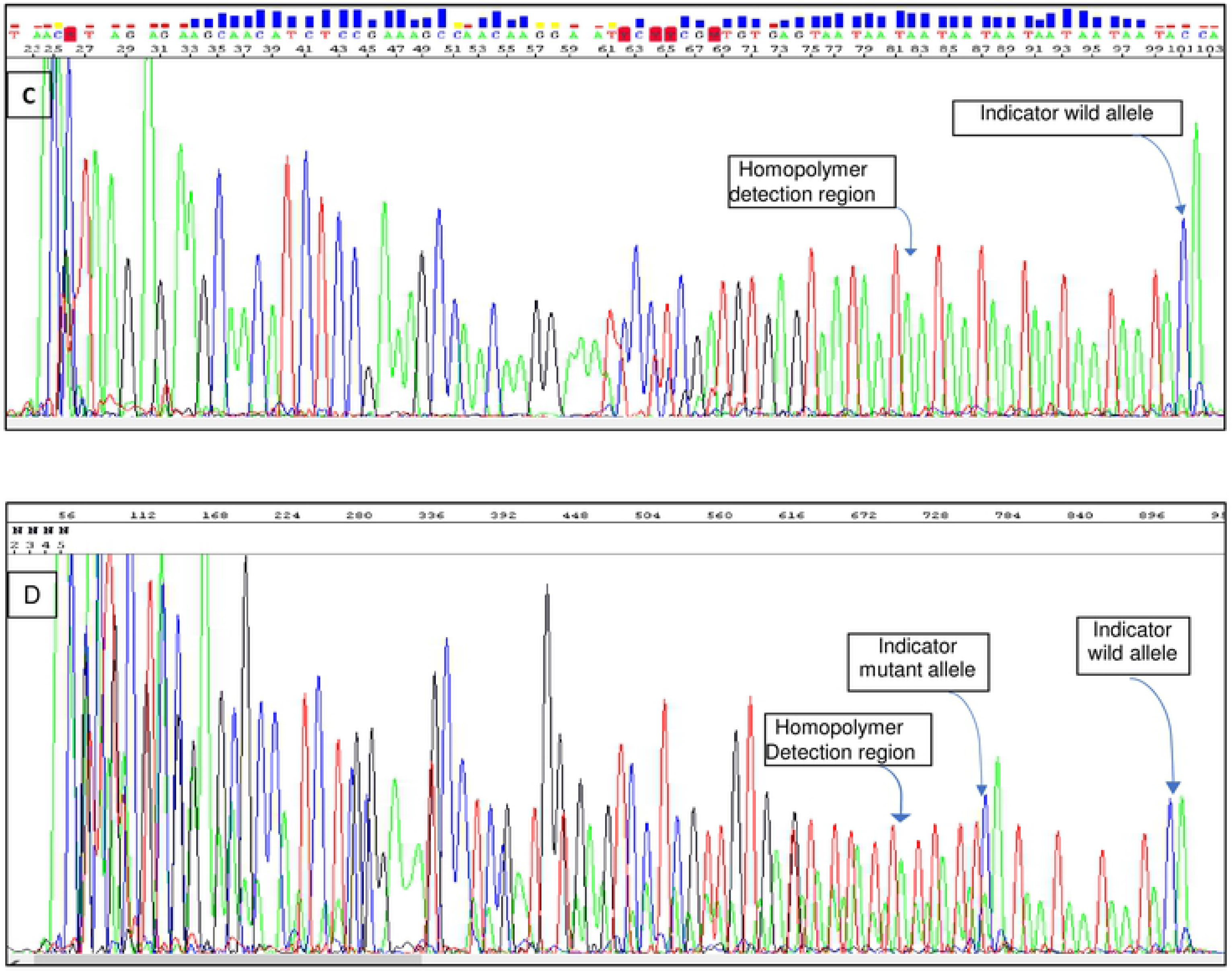
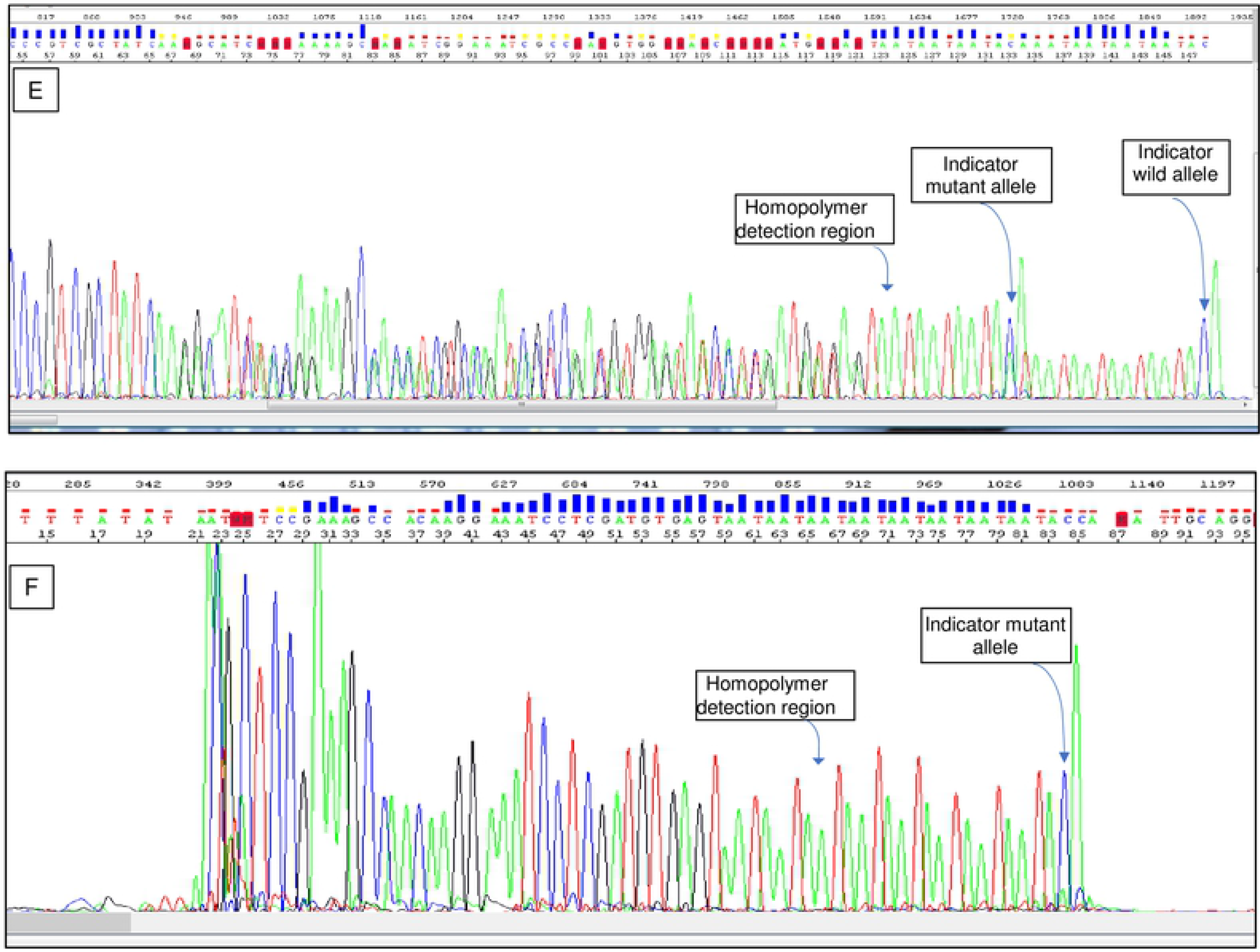
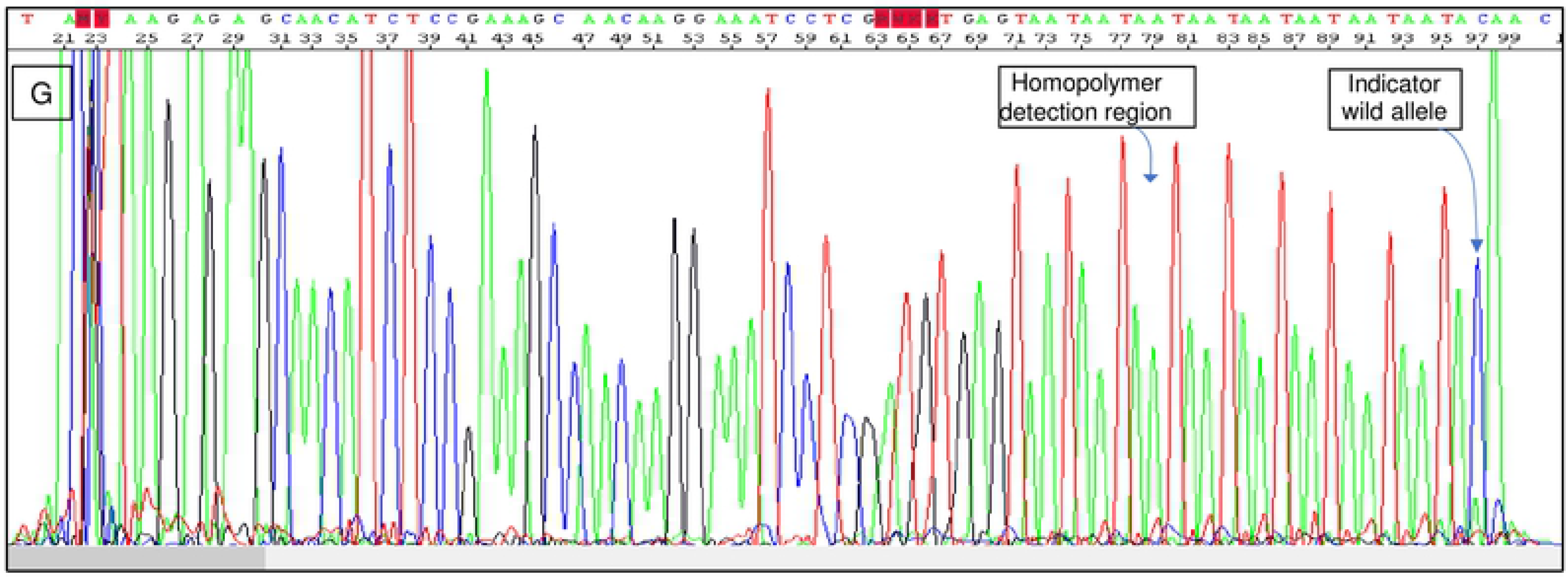
RepSeq technology design and sequencing result. Panel A: Diagrammatic representation of various elements of RepSeq technology. Panel B: Showing the location of sequencing primers, deletion region lower primer and detection region. Panel C: Electropherogram generated using wild type sequencing primer (selective) with homozygous DNA template HD 709. Panel D: Electropherogram generated using wild type sequencing primer (selective) and deletion sequencing primers with heterozygous DNA template 251 carrying 50% deletion. Panel E: Electropherogram generated using wild type sequencing primer (non -selective) with heterozygous DNA template 251. Panel F: Electropherogram generated using deletion sequencing primer with DNA extracted from human cell line HD 251 carrying 50% deletion. Panel G: Electropherogram generated using both deletion sequencing primer and wild type sequencing primers (selective) with homozygous DNA template HD 709.

#### Samples

The study included four categories of samples; DNA extracts from human cell lines obtained from Horizon Discoveries (Cambridge, UK), twenty-one de-identified blood samples from normal individuals, twenty-four de-identified DNA extracts that were previously tested by pyrosequencing, and twenty de-identified DNA extracts that were previously tested by TruSeq cancer panel. This manuscript is focused on technic method. All the samples have been de-identified. Therefore, IRB approval is not needed.

#### Sample Preparation

Total DNA was extracted from 1 ml of blood sample using DNeasy Blood and Tissue Kit (Qiagen, USA). The concentration of DNA ranged from 2.2 ng/ul to 9.7 ng/ul, and 10 ul of the DNA extract was used per reaction. DNA concentration of samples that were previously tested by TruSeq cancer panel ranged from 10ng/ul to 139 ng/ul. The samples were diluted 1:3 in TE buffer and 10 ul of the diluent was used in the PCR reaction. DNA concentration of samples that were previously tested by Pyrosequencing was not determined. Ten microliters of the diluent were used in the PCR reaction.

#### Amplification

Two step PCR Each PCR reaction included 25.0 μl of 2X buffer (MultiPlex PCR Master Mix, Qiagen). One microliter each of the 10 pmol forward and reverse primers (Select MultiGEN Diagnostics, USA) were added to the reaction with 10.0 μl of the sample DNA extract. The first thermocycling was carried out at: 95°C/5min, (95°C/30sec, 57.5°C/90sec, 72°C/30sec) x 20 cycles, 25°C/10min. Twenty-five microliters of Buffer 2 (Select MultiGEN Diagnostics Inc, USA) was added and second thermocycling with the above conditions was performed with 20 cycles. Following PCR clean-up with Ampure (Beckman, Agencourt USA), a sequencing reaction was set up using 1.0 μl of Big dye, 9.5 μl of 5X sequencing buffer, 1 pmol of sequencing primer (Select MultiGEN Diagnostics, USA), 30 μl of the purified PCR products and 4.5 μl of Dnase free water to a total reaction volume of 50 μl. Cycle Sequencing conditions: (96°C/105sec, 55°C/10sec, 60°C/2.5min) x 25 cycles. Cycle sequencing products were cleaned with CleanSEQ (Beckman Agencourt USA) and eluted in 40 μl of Dnase free water. The cleaned products were injected for 16 seconds into ABI 3130xl Genetic Analyzer and the electropherogram was analyzed using Sequencing Analysis Software 6.0.

### Result analysis

If there is no deletion, and using selective sequencing primer only, the single nucleotide sequence of the wild type will be displayed in the sequencing result with its specific read sequences (Figure 1C). If the sample carries cells with EGFR deletion, then there will be two nucleotide sequences; one from the wild-type sequence and other from that of EGFR deletion, and hence the nucleotide signal from both wild type and mutant will overlap for most part. Although at the beginning of the electropherogram the overlap of both the nucleotide sequences could be scrambled and not readable, the nucleotide sequences at the distal end will display the detection region that is made up of adenosine thymidine-thymidine repetitive homopolymer sequence with cytosine at its distal end (Figure 1D). Since the nucleotide sequence from the deletion will be shorter, the indicator cytosine residue on the deletion sequence will move to the left on the electropherogram and will be in the midst of the thymidine-adenosine-adenosine repetitive homopolymer sequence detection region, thus the presence of deletion can be easily recognized and therefore confirmed.

### Limit of detection

Limit of detection was determined using a known amount of human gDNA extracted from human cell line (HD 251). Stock solution of 5ug/ul was 1 in 10 serially diluted. Five microliters of diluted stock were used in a PCR reaction and the lower limit of detection was determined to be 1.47 copies per assay. (Table 1).

**Table 1.** Lower limit of detection.

| Serial dilution | Copies/Rx (5ul) | Results |
| --- | --- | --- |
| Stock solution | 147.2 | Positive for wild and L747_A750 |
| 1:10 | 14.7 | Positive for wild and L747_A750 |
| 1:100 | 1.47 | Positive for wild and L747_A750 |
| 1:1000 | <1.0 | Negative for wild and L747_A750 |
| 1:10000 | <1.0 | Negative for wild and L747_A750 |
| 1:100000 | <1.0 | Negative for wild and L747_A750 |

Note: Stock solution 5ng/ul

## Results and Discussion

### Verification of Assay specificity

The specificities of the deletion sequencing primer and the wild type sequencing primer were cross-checked using two DNA templates extracted from human cell lines (Horizon Discoveries Inc): HD 251 carrying 5 0 % deletion and 50% wild type DNA, and HD 709 carrying 100% wild type DNA. Various combinations of the sequencing primers and the templates were tested (Table 2). The wild type sequencing primer (selective) generated a wild type nucleotide sequence when tested with wild type template HD 709 (Figure 1C). The wild type sequencing primer (Non-selective) generated a wild type nucleotide sequence and deletion sequence with heterozygous template (HD 251) that carries 50% wild type allele (Figure 1E). The deletion sequencing primer generated a deletion specific nucleotide sequence when tested with heterozygous template (HD 251) that carries 50% wild type allele (Figure 1F). However, the mutant sequencing primer did not generate any nucleotide sequence when tested with 100% wild type template HD 709 (Data not shown). Further, when mutant and wildtype (selective) sequencing primers were both included with wild type template (HD 709), only wild type specific and no deletion specific sequences were generated (Figure 1G). When both the wild type (selective) and the deletion sequencing primers were tested with heterozygous template (HD 251), the expected nucleotide sequences were generated, with the indicator signal for the deletion moving to the left of the wild type (Figure 1D).

**Table 2.** Primer specificity.

| Template |  | Sequencing primer<br>(Selective) | Electropherogram |
| --- | --- | --- | --- |
| Name | Composition |  |  |
| HD251 | Mutant:Wild=50%:50% | Mutant | Mutant sequence |
| HD709 | Wild=100% | Mutant | Negative- No sequence |
| HD251 | Mutant:Wild=50%:50% | Wild | Wild sequence |
| HD709 | Wild=100% | Wild | Wild sequence |
| HD251 | Mutant:Wild=50%:50% | Mutant /Wild | Mutant & Wild sequence |
| HD709 | Wild=100% | Mutant /Wild | Wild sequence |

Note: HD Horizon Discoveries, UK

The human genomic controls with 50% mutant allele template generated both wild type sequence and the deletion sequence. In addition, twenty-one normal blood samples were tested for EGFR deletion L747_A750 by RepSeq technology using both deletion and wild type specific sequencing primers, and the results from all twenty-one samples generated only wild type sequence and no sequence indicative of deletion.

### Comparison with pyrosequencing

Out of the twenty-four samples that were tested by pyrosequencing; both methods detected 15 EGFR L747-A750 negatives and nine EGFR L747-A750 positives. (Table 3).

**Table 3.**
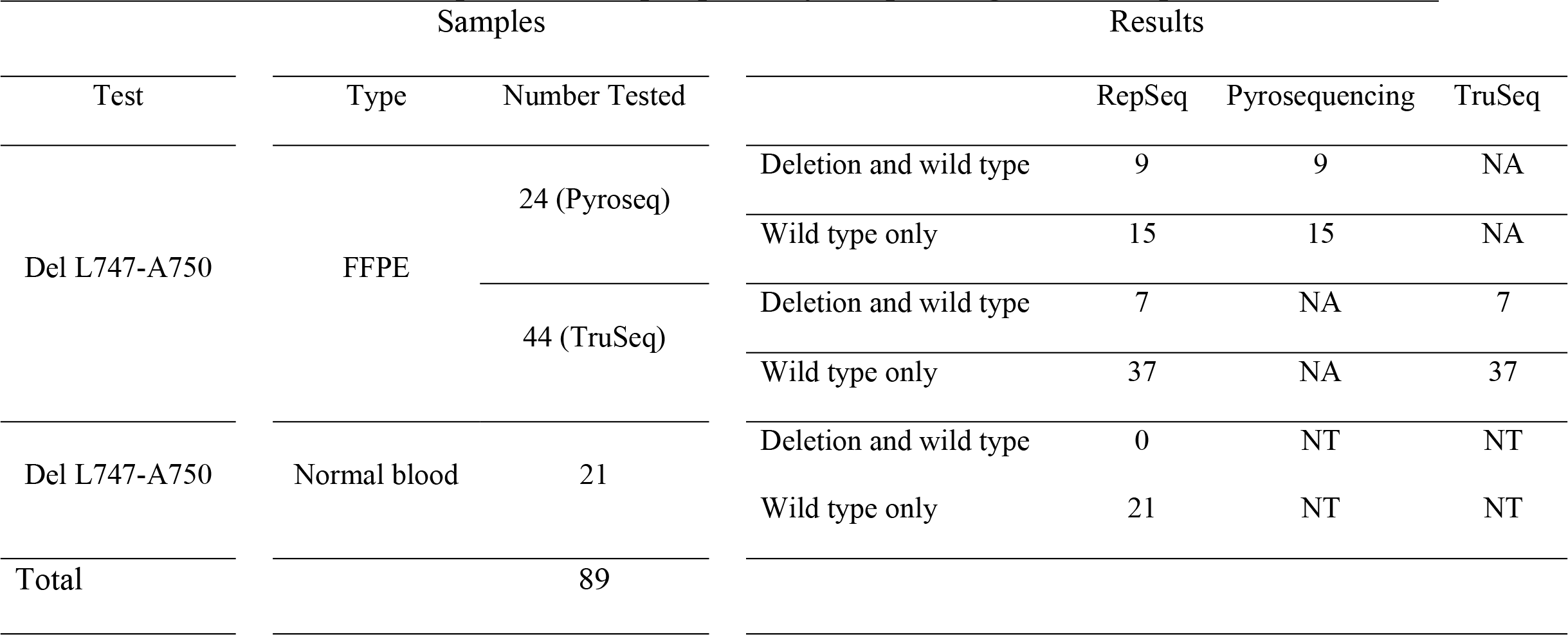
Comparison of RepSeq with Pyrosequencing and Truseq.

### Comparison with TruSeq

Out of forty-four FFPE samples tested, RepSeq detected all seven EGFR L747-A750 positives and thirty-seven EGFR L747-A750 negatives that were detected by TruSeq negatives. (Table 3).

### RepSeq features

Similar to Sanger sequencing and pyrosequencing, the electropherogram from the wild type and the deletion are generated from the same set of PCR primers. However, unlike Sanger sequencing and pyrosequencing, RepSeq uses two sequencing primers; one for the wild type and the other for a deletion-specific sequencing primer that spans across the deletion region. This unique feature increases the signal intensity of that of the deletion that is in par with that of the wild type and therefore increases the test sensitivity. The lower PCR primer carries a three nucleotide (Adenosine-Thymidine-Thymidine) repetitive sequence with a guanidine nucleotide at its 5’ end as an indicator of one end of the PCR product. Such a design make sequencing data interpretation much easier and faster.

This study used FFPE samples from late stage lung cancer where positive EGFR L747_A750 deletion samples will have an abundance of copies of the deletion, and hence had acceptable level of concordance among the three methodologies tested. Since RepSeq has a very low limit of detection compared to other two methods, it is expected to play a significant role in liquid biopsy and detection of EGFR 747_A750 in early stages (< stage IV).

There are other additional deletions that are clinically significant. A second generation RepSeq-EGFR assay will have a combination of sequencing primers covering additional deletions. The RepSeq platform also could be applied to other clinically significant variations, such as ALK-EML4, ROS (9).

In summary, the EGFR RepSeq assay produces an easy to read electropherogram to detect mutation in the presence of wild type, and enrichment using allele specific primers. Further, RepSeq also contains built-in features that address troubleshooting due to variations in sample matrix.

